# Occurrence of *Cryptosporidium hominis* in cattle bordering the Lake Mburo National Park in Kiruhura district, Western Uganda

**DOI:** 10.1101/562793

**Authors:** Sarah Gift Witto, Clovice Kankya, Anne Kazibwe, Gloria Akurut, Sylvester Ochwo

**Affiliations:** Molecular Biology Laboratory, Department of Biomolecular Resources and Biolab Sciences, College of Veterinary Medicine Animal Resources and Biosecurity, Makerere University, P. O. Box 7062 Kampala, Uganda; Department of Biosecurity, Ecosystems and Veterinary Public Health, College of Veterinary Medicine Animal Resources and Biosecurity, Makerere University, P. O. Box 7062 Kampala, Uganda; Department of Biomolecular Resources and Biolab Sciences, College of Veterinary Medicine Animal Resources and Biosecurity, Makerere University, P. O. Box 7062 Kampala, Uganda

**Keywords:** *Cryptosporidium hominis*, Cattle, Polymerase chain reaction, Genotyping, Uganda

## Abstract

**Background:** *Cryptosporidium* is an emerging opportunistic zoonotic pathogen that causes diarrheal illness in a wide range of hosts including livestock and humans. Globally there is exponential increase in livestock production to meet the worlds’ demand for animal protein as well as for financial reasons. However, there is raised concern of the public health threat due to contamination of the environment by livestock waste carrying zoonotic pathogens such as *Cryptosporidium*. This study set out to establish the prevalence of *Cryptosporidium* as well as the circulating genotypes in order to elucidate the potential role of cattle in the spread of human cryptosporidiosis. We collected rectal coprological samples from 363 cattle in 11 households in Kiruhura district, Southwestern Uganda. The samples were screened for presence of *Cryptosporidium* oocysts using the phenol auramine staining method followed by fluorescent microscopy. DNA was then extracted from the microscopy positive samples and the COWP gene amplified using PCR. Amplified gene products were sequenced and subjected to phylogenetic analysis.

**Results:** The overall animal level prevalence of *Cryptosporidium* was 7.7% (95% CI: 5.1-10.9), and herd level prevalence was 33.3% (95% CI: 18.5-52.2). We found a statistically significant difference (p=0.02) between infection in bulls as compared to cows. There was however no significant difference in the prevalence among the different cattle breeds sampled, with the following prevalence’s observed in Crosses 9.2%, Ankole 5.7%, Friesian 7.1%, and Boran 2.8% respectively. The COWP gene was successfully amplified from 20 of the 28 microscopy positive samples. All the sequenced DNA amplicons were confirmed to be *C. hominis*, with 98%-100% identity to sequences in the GenBank. *C. hominis* was the only genotype isolated from this study, further asserting that cattle could be a potential high risk source of human cryptosporidiosis.

**Conclusion:** This study represents the first time naturally occurring *C. hominis* has been isolated from cattle in Uganda. This further provides evidence of cattle possibly being biological reservoirs for *C. hominis* and cattle could be a potential high risk source of human cryptosporidiosis.

## Background

*Cryptosporidium* is an emerging zoonotic enteric pathogen that causes diarrheal illness in both humans and animals known as cryptosporidiosis (Guerrant, 1997). Cryptosporidiosis infections in cattle are more prevalent in calves as compared to the adult animals; clinical signs occur 3-5 days after infection and include profuse watery diarrhea, gastrointestinal discomfort, nausea as well as fever (De Graaf, Vanopdenbosch, Ortega-Mora, Abbassi, & Peeters, 1999; Fiuza *et al*., 2011). These episodes normally result into weight loss and occasionally death (Rajendran *et al*., 2011; Ryan *et al*., 2005).

Twenty six species of *Cryptosporidium* are currently documented (Ryan, Fayer, & Xiao, 2014) and they infect a wide range of animal species. The important *Cryptosporidium* species which infect cattle are *C. parvum, C. bovis*, and *C. andersoni* (Fayer, Santin, & Trout, 2007) however, other *Cryptosporidium* species and genotypes have sporadically been reported in cattle but these lack epidemiological significance (Fiuza *et al*., 2011; Ralston, 2009). Cattle are the biological reservoir for *C. parvum*, a zoonotic species commonly implicated in outbreaks of human cryptosporidiosis (Blackburn *et al*., 2006; Millard, P. S., Gensheimer, K. F., Addiss, D. G., Sosin, D. M., Beckett, G. A., Houck-Jankoski, A., & Hudson, 1994; Slifko, Smith, & Rose, 2000).

In humans, *C. hominis* is the main cause of disease and is considered host specific however recent studies have reported isolation of *C. hominis* in livestock (Rajendran *et al*., 2011; Smith *et al*., 2005; Xiao & Fayer, 2008). The arthroponotic transmission of *C. hominis* (environmental loading of wastes) is a public health concern especially in Sub Saharan Africa because almost a quarter of the people lack access to safe drinking water and basic sanitation which are risk factors. The poor sanitation and lack of safe drinking water coupled with the HIV burden has resulted in an enhanced burden of human cryptosporidiosis (Aldeyarbi, Abu El-Ezz, & Karanis, 2016).

The global burden of human cryptosporidiosis is unknown however according to Kotloff *et al*., (2013) *Cryptosporidium* is one of the leading causes of moderate-to-severe diarrhea and the second most common pathogen in children living in Sub-Saharan Africa. Diarrheal episodes in Sub Saharan Africa is reported to be responsible of 14% hospital outpatient visits, 16% of hospital admissions and an average of 35 days of illness per year in children (Greenwood *et al*., 1987) and an estimated 1.8 million deaths (Wardlaw, Salama, Brocklehurst, Chopra, & Mason, 2010). In non-fatal cases of diarrhea, particularly chronic infections diarrhea has been strongly correlated with growth retardation and yet good health is a precondition for society to develop. (Prado *et al*., 2005; Thompson, 2008; Tumwine *et al*., 2003)

Several studies continue to elucidate the role played by livestock in the transmission of *Cryptosporidium* to humans (Giles *et al*., 2009; Gormley, Little, Chalmers, Rawal, & Adak, 2011; Kang’ethe *et al*., 2012; Rajendran *et al*., 2011; Samra, 2013). These studies provide an in-depth understanding of the host range of *Cryptosporidium* which is crucial in the development of strategies that prevent both the anthroponotic and the zoonotic transmission of the disease (Giles *et al*., 2009; Rajendran *et al*., 2011). *Cryptosporidium* is transmitted via the fecal oral route through the ingestion of water or food contaminated with oocysts. Oocysts may also be ingested through direct contact with fecal material from individuals (Blackburn *et al*., 2006; Ponka *et al*., 2009; Slifko *et al*., 2000). *Cryptosporidium* has a low infective dose with as few as 9 oocysts capable of causing disease (Okhuysen, Chappell, & Crabb, 1999).

In this study we report the occurrence of *C. hominis* in cattle from south western Uganda. These are communities where the water sources are shared amongst livestock and humans. This information we hope will contribute more knowledge about the epidemiology of Cryptosporidiosis and further contribute to the formulation of control strategies to protect high-risk populations from disease resulting from either human or animal hosts.

## Methods

### Study area

Nyakashashara and Sanga sub-counties are found in Kiruhura district located in the Western Region of Uganda bordering Lake Mburo National Park (LMNP) (Fig.1). Kiruhura is a water stressed area characterised by drought conditions with scarce potential for ground water. It has a human population of 280,200 and 81% of households use open water sources (Kiruhura District Local Government, 2012). Kiruhura is a farming district with a cattle population of 342,315 (MAAIF & UBOS, 2015). Livestock forms the backbone of economic activity in the district.

**Figure 1:**
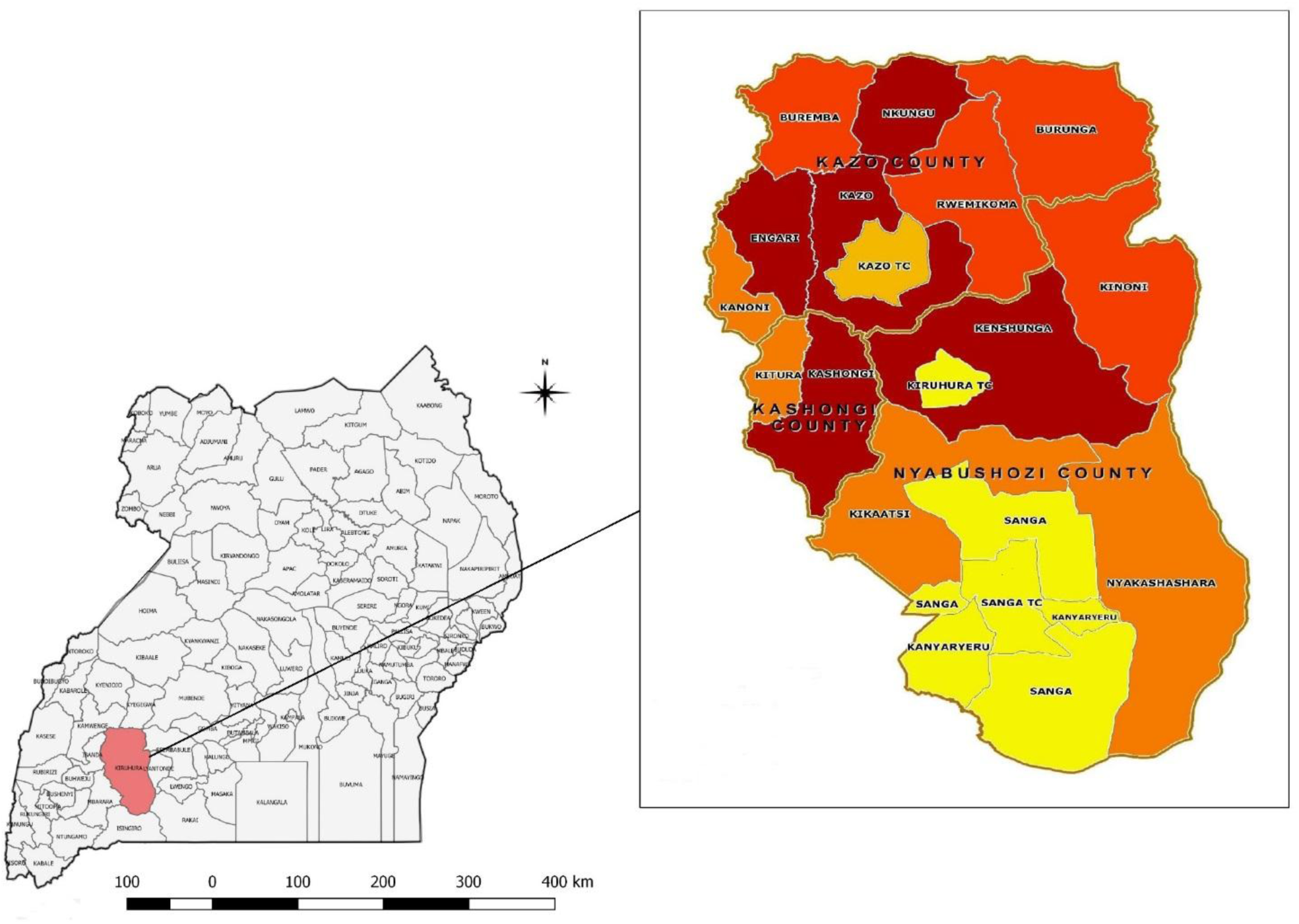
Map of Uganda showing location of study area.

### Study design

This was a cross-sectional study done in February to March 2014. Cattle were sampled from 11 farms in Nyakashashara and Sanga Sub-counties in Kiruhura district, Western Uganda. Farms with unprotected water sources, shared by both humans and livestock were selected. Within the selected farms, simple random sampling was used to select cattle to be sampled.

### Sample size determination

This cross-sectional study was conducted between February 2014 and March 2014 in Nyakashashara and Sanga Sub-counties in Kiruhura district, Western Uganda. The sample size was determined by the Kish and Leslie formula for cross-sectional studies. A prevalence of 38% for *Cryptosporidium* (Nizeyi, Cranfield, & Graczyk, 2002) was used to calculate the sample size N = (Z^2*^P) (1-P)/d^2^

Where N is the sample size, Z^2^ is the abscissa of the normal curve that cuts off an area at 1.96 (1-equals the desired confidence level, e.g., 95%), d is the desired level of precision of 0.05, P is the estimated proportion of an attribute that is present in the population of 0.38 for *Cryptosporidium*. Therefore; N= (1.96^2^*0.38) (1-0.38)/ 0.05^2^

N= 362, 363 fecal samples were collected and examined.

### Sample collection

Rectal fecal specimens from 363 cattle, each weighing approximately 10g were collected from eleven (11) farms located in Nyakashashara and Sanga sub-counties. Each specimen was placed into a sterile container and sealed. Details of location, age and sex of animals were recorded and the specimens were transported in a cool box at 4°C to the Molecular Biology Laboratory, Makerere University for analysis.

### Formalin diethyl ether concentration

Approximately 3g of the fecal samples were individually weighed and homogenized with 3ml of phosphate-buffered saline (1Χ PBS) pH 7.4 (Nizeyi *et al*., 2002). The homogenate was sieved with cotton gauze and transferred to 15 ml falcon tube. After sieving the homogenate, 7ml of 10% formalin and 3 ml of diethyl ether were added, hand shaken and the mixture was centrifuged at 2000 rpm for 3 minutes. The diethyl ether layer, the particulate plug and the formalin below it were discarded and the sediment was retained for examination (Alexander, 2014).

### Auramine-phenol staining and microscopic analysis

The sediment was washed in 10ml of 1Χ PBS) pH 7.4 and span at 5000g for 10 minutes and the supernatant discarded. This process was repeated three times. The sediment was re-suspended in 200µl of 1Χ PBS pH 7.4 and 50µl of the mixture was used to prepare smears on slides. The slides were air dried and fixed with absolute methanol for 3 minutes before staining.

The slides were stained using the auramine phenol technique according to the Alexander (2014). The slides were immersed in auramine phenol stain for 10 minutes. The stain was then rinsed off in tap water and the smears decolorized with 3% acid alcohol for 5 minutes. The smears were counterstained in 0.1% potassium permanganate for 30 seconds and rinsed in water to remove the excess stain. The smears were air dried at room temperature and examined for the presence of oocysts, using a fluorescence microscope equipped with FITC filters, by scanning the slide under the ×20 objective lens and confirming for the presence of oocysts under the ×40 objective lens.

### DNA extraction

DNA was extracted from 150µl of each re-suspended fecal sediment, using the QIAamp Fast DNA Stool Mini Kit (QIAGEN, Hilden, Germany). The extracted DNA was stored at −20 °C for Polymerase Chain Reaction (PCR).

### PCR amplification *Cryptosporidium* COWP gene

DNA extracted from the oocysts was used to amplify the 553bp fragment of the COWP gene using a nested PCR (Spano, Putignani, McLauchlin, Casemore, & Crisanti, 1997). PCR amplification was performed in 25µl volumes with 2Χ Ready-mix (Bioline, UK) (volume 12.5µl, final concentration 1Χ), forward primer (CWPF), 10pmol (volume 2.5µl, final concentration 1.0µm), reverse primer (CWPR), 10pmol (volume 2.5µl, final concentration 1.0µm), DNA template 2.5µl, nuclease free water to 25µl. A PCR mastermix without template DNA was used as a negative control and included in each experiment. A positive control was also included. The following cycling conditions were used; initial denaturation for 5 minutes at 94°C, followed by 50 cycles of denaturation at 94°C for 30 seconds, annealing 55°C for 1 minute and extension 72°C for 45 seconds with a final extension of 72°C for 7 min. and a 12°C hold. A second run was performed on the samples with the second set of primers (Cry9 and Cry 15). The PCR conditions were identical to the one in the primary run except the annealing temperature which was reduced to 52°C for 1 minute. PCR products were separated on a 1% agarose gel stained with ethidium bromide and visualized using a UV gel documentation system (UV pro). A 1kb molecular weight marker (Invitrogen®) was used during the agarose electrophoresis as a standard. All the nested PCR products of COWP genes were purified using a DNA purification kit (QIAGEN, Germany). The quality and quantity of the purified PCR products was checked with the NanoDrop 1000 spectrophotometer (Thermo Fisher, USA) and then sent for Sanger sequencing at Inqaba biotech in South Africa.

**Table 1:**
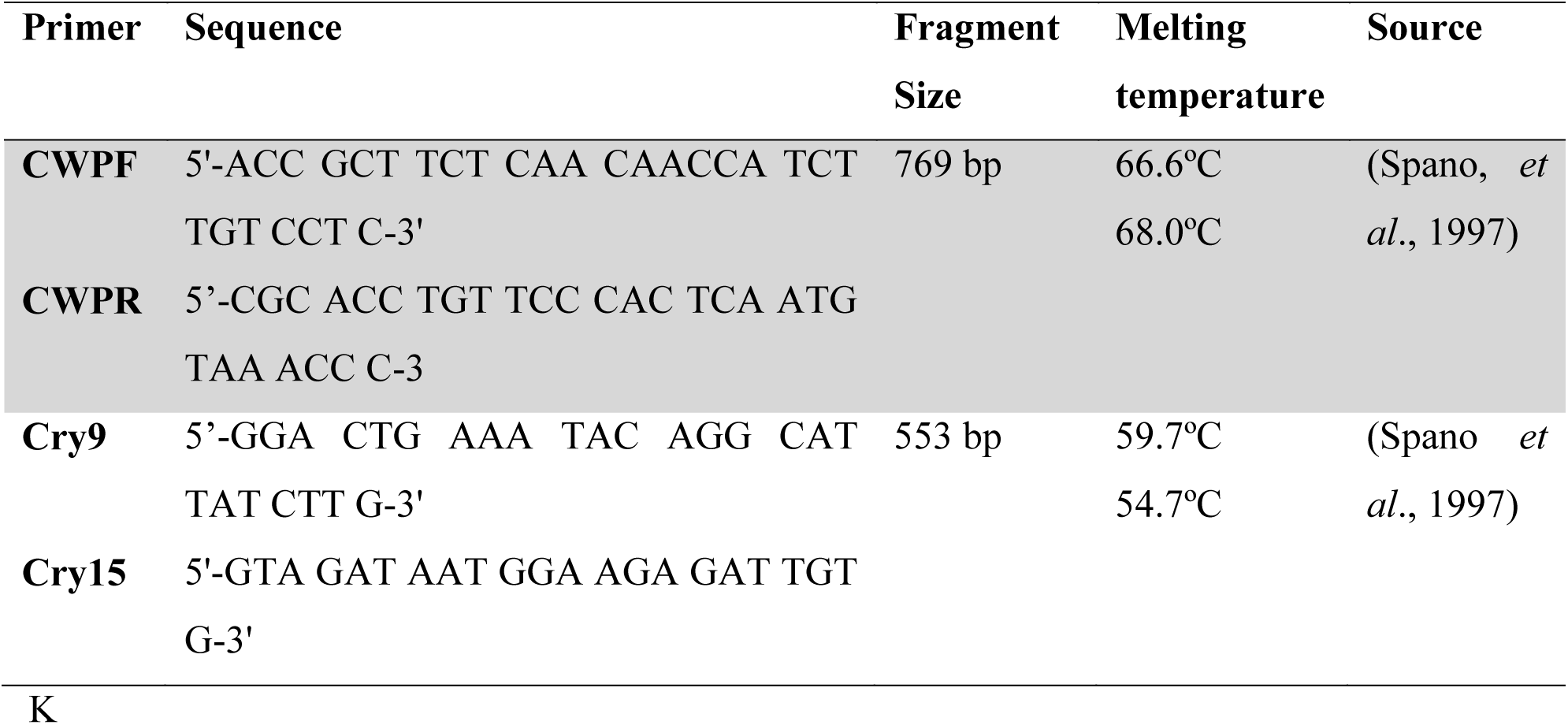
COWP primer sequences

### DNA Sequencing

DNA Sequencing was done by a commercial company (Inqaba biotech, South Africa), using the Sanger sequencing method.

### Analysis of COWP gene sequences

To determine the taxonomic positions of newly generated COWP sequences relative to published sequences, phylogenetic trees were constructed using the Maximum Likelihood method based on the Tamura 3-parameter model in the computer program MEGA6. The robustness of groupings was assessed using 1000 bootstrap replicates of the data (Tamura, Stecher, Peterson, Filipski, & Kumar, 2017). All sequences generated during this study were deposited in GenBank and assigned accession numbers KY586953-KY586963.

## Results

### Prevalence of *Cryptosporidium* as quantified by Microscopy

The overall prevalence of *Cryptosporidium* infections in the cattle quantified by microscopy using phenol auramine staining method was 7.7% (28/ 363). Farm 2 had the highest infection rate (33.3%), followed by farm 7 (25%), 8 (18.5%), 11 (7.1%), 1 (6.9%), 6 (4.3%) and lastly 4 (3.6%).

**Table 2:**
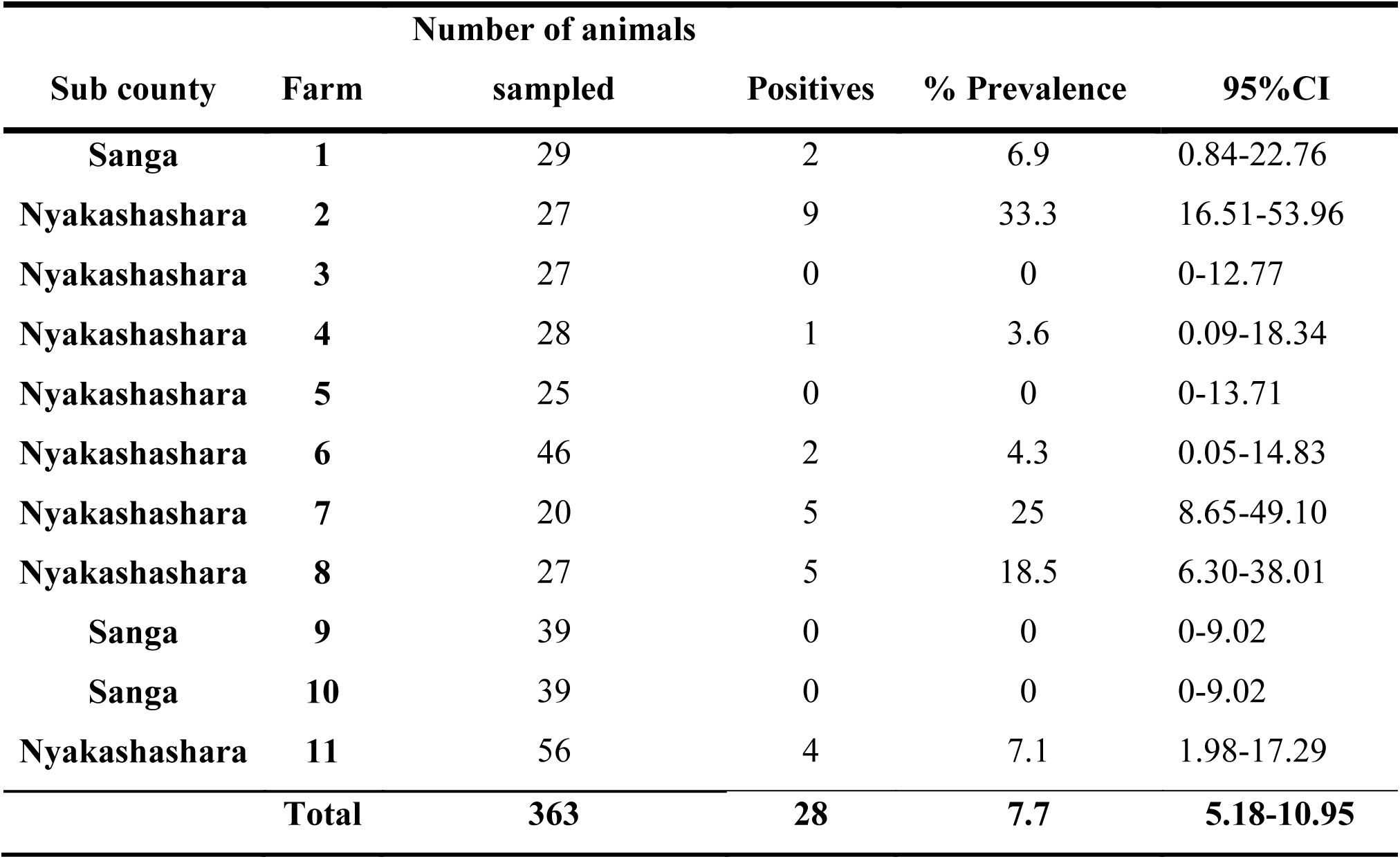
Prevalence of *Cryptosporidium* species in cattle in the study sites

| Number of animals |  |  |  |  |  |
| --- | --- | --- | --- | --- | --- |
| Sub county | Farm | sampld | Positives | % Prevalence | 95%CI |
| **Sanga** | **1** | 29 | 2 | 6.9 | 0.84-22.76 |
| **Nyakashashara** | **2** | 27 | 9 | 33.3 | 16.51-53.96 |
| **Nyakashashara** | **3** | 27 | 0 | 0 | 0-12.77 |
| **Nyakashashara** | **4** | 28 | 1 | 3.6 | 0.09-18.34 |
| **Nyakashashara** | **5** | 25 | 0 | 0 | 0-13.71 |
| **Nyakashashara** | **6** | 46 | 2 | 4.3 | 0.05-14.83 |
| **Nyakashashara** | **7** | 20 | 5 | 25 | 8.65-49.10 |
| **Nyakashashara** | **8** | 27 | 5 | 18.5 | 6.30-38.01 |
| **Sanga** | **9** | 39 | 0 | 0 | 0-9.02 |
| **Sanga** | **10** | 39 | 0 | 0 | 0-9.02 |
| **Nyakashashara** | **11** | 56 | 4 | 7.1 | 1.98-17.29 |
| **Total** |  | **363** | **28** | **7.7** | **5.18-10.95** |

A difference in the prevalence by breed was observed (Crosses 9.2%, Ankole 5.7%, Friesian 7.1%, Boran 2.8%), (Table 3). However the difference observed was found to be statistically insignificant (Table 4).

**Table 3:** Prevalence of *Cryptosporidium* by breed

|  | Number of Positive | Number sampled | % Prevalence | (95% CI) |
| --- | --- | --- | --- | --- |
| **BORAN** | 1 | 36 | 2.8 | 0.07-14.52 |
| **ANKOLE** | 3 | 53 | 5.7 | 1.18-15.66 |
| **CROSSES** | 20 | 218 | 9.2 | 5.69-13.81 |
| **FRESIAN** | 4 | 56 | 7.1 | 1.98-17.29 |
| **TOTAL** | **28** | **363** | **7.7** | **5.18-10.95** |

**Table 4:** Risk factors for *Cryptosporidium* infection

| Variable | OR (Odds ratio) | p-value |
| --- | --- | --- |
| **Sex** |  |  |
| Cow | 0.17 | **0.022†** |
| Bull | 1.00 |  |
| **Breed** |  |  |
| Ankole | 2.14 | 0.52 |
| Friesian | 2.69 | 0.39 |
| Cross | 3.57 | 0.22 |
| Boran (reference) | 1.00 |  |
| **Age** |  |  |
| Calves | 1.34 | 0.802 |
| Yearling | 0 |  |
| Heifers (reference) | 1.00 |  |

† Significant p-value

### Risk factors for infection with *Cryptosporidium*

Risk factor analysis of breed, age and sex showed sex as a risk factor. Cows are 83% less likely of contracting *Cryptosporidium* than the bulls. However differences in prevalence by age and breed were not statistically significant as shown in table 4.

### PCR amplification of the COWP gene

Genomic DNA was extracted from the 28 microscopy positives samples. Of the 28 positive samples, the 553bp COWP gene product was successfully amplified in 20 samples. Failure to amplify the COWP gene product in the 8 samples could be due to fecal constituents such as bilirubin, bile salts, and complex polysaccharides which inhibit PCR even when present at low concentrations (Morgan, F. U., Pallant, L., Dwyer, B. W., Forbes, D. A., Rich, G., & Thompson, 1998; Thornton & Passen, 2004).

**Figure 2:**
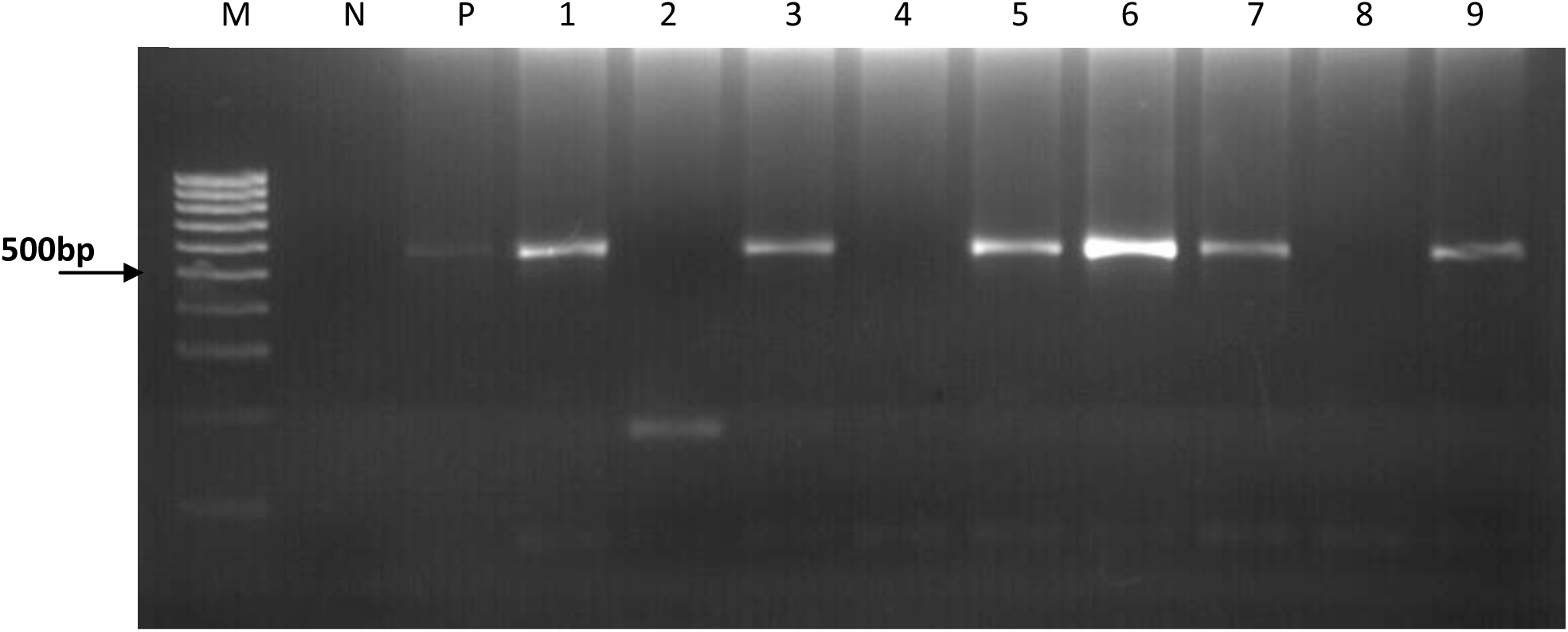
A representative 2% agarose gel showing the amplification of the 553bp fragment of the COWP gene. Lane M is a 1kb molecular weight marker, lane N is the negative control and lane P is the positive control. Lanes 1, 3, 5, 6, 7 and 9 are positive samples with a 553bp band size. Lanes 2, 4 and 8 are negative samples.

### Sequence analysis of *Cryptosporidium* COWP gene

The PCR products of the 553bp COWP gene amplification from 11 of the 20 samples were successfully sequenced by Sanger method. All the sequences were identified as *C. hominis* by BLAST search and had 98%-100% identity to sequences in the GenBank as shown in table 5.

**Table 5:**
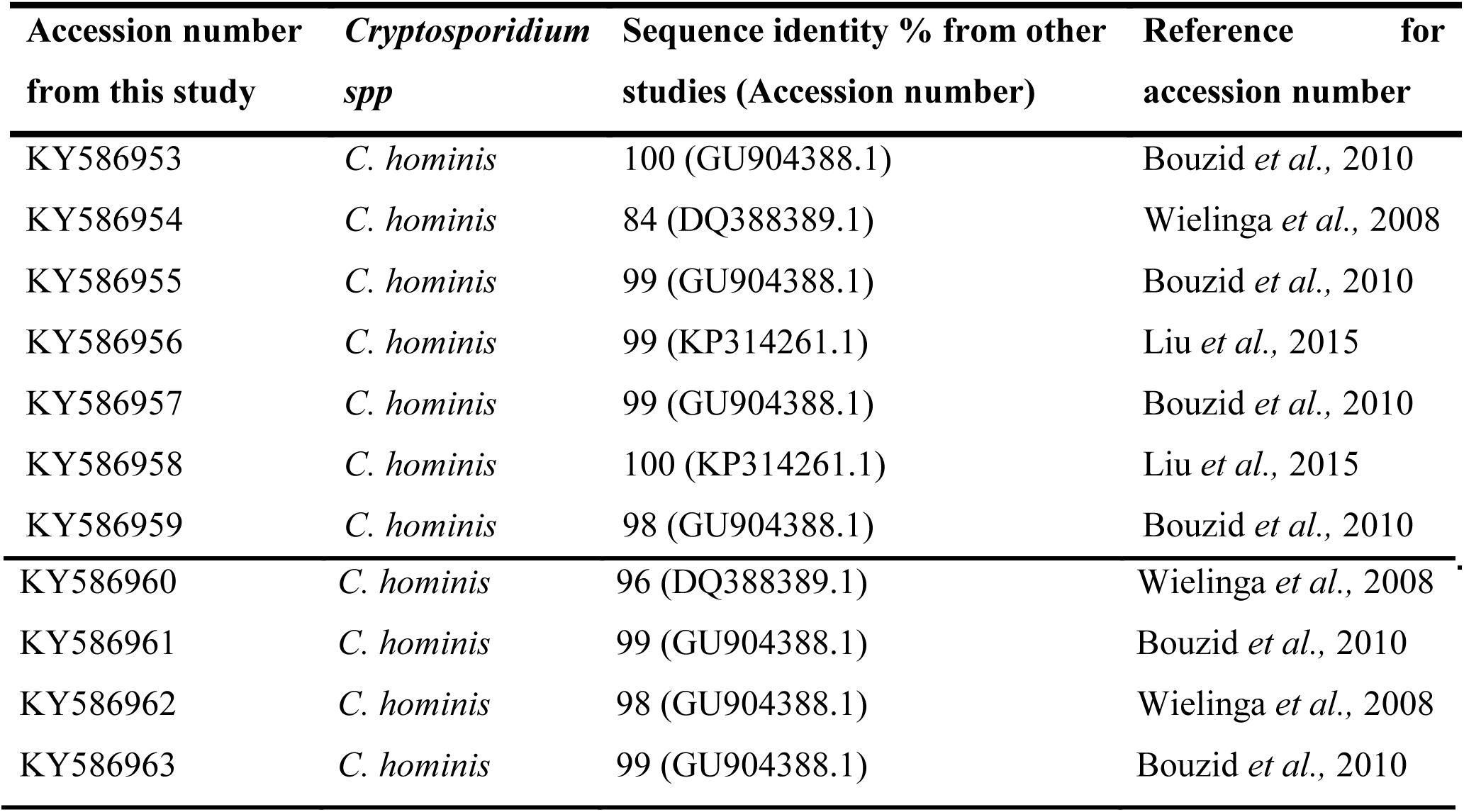
*Cryptosporidium* species detected by PCR and sequencing of the COWP gene in faecal samples collected from in Kiruhura District, Uganda

### Phylogenetic analysis of the COWP gene

The nucleotide sequences of the COWP gene fragment were aligned using ClustalW (see additional file 1) and showed that the sequences from this study were highly identical to *C. hominis* sequences from the GenBank. Phylogenetic analysis of the newly generated COWP gene sequences and representative published sequences yielded a tree where all *Cryptosporidium* sequences from this study (KY586953-KY586963) clustered within a clade containing known *C. hominis* sequences with a bootstrap value of 95, therefore indicating that this clade is highly supported (Figure 3)

**Figure 3:**
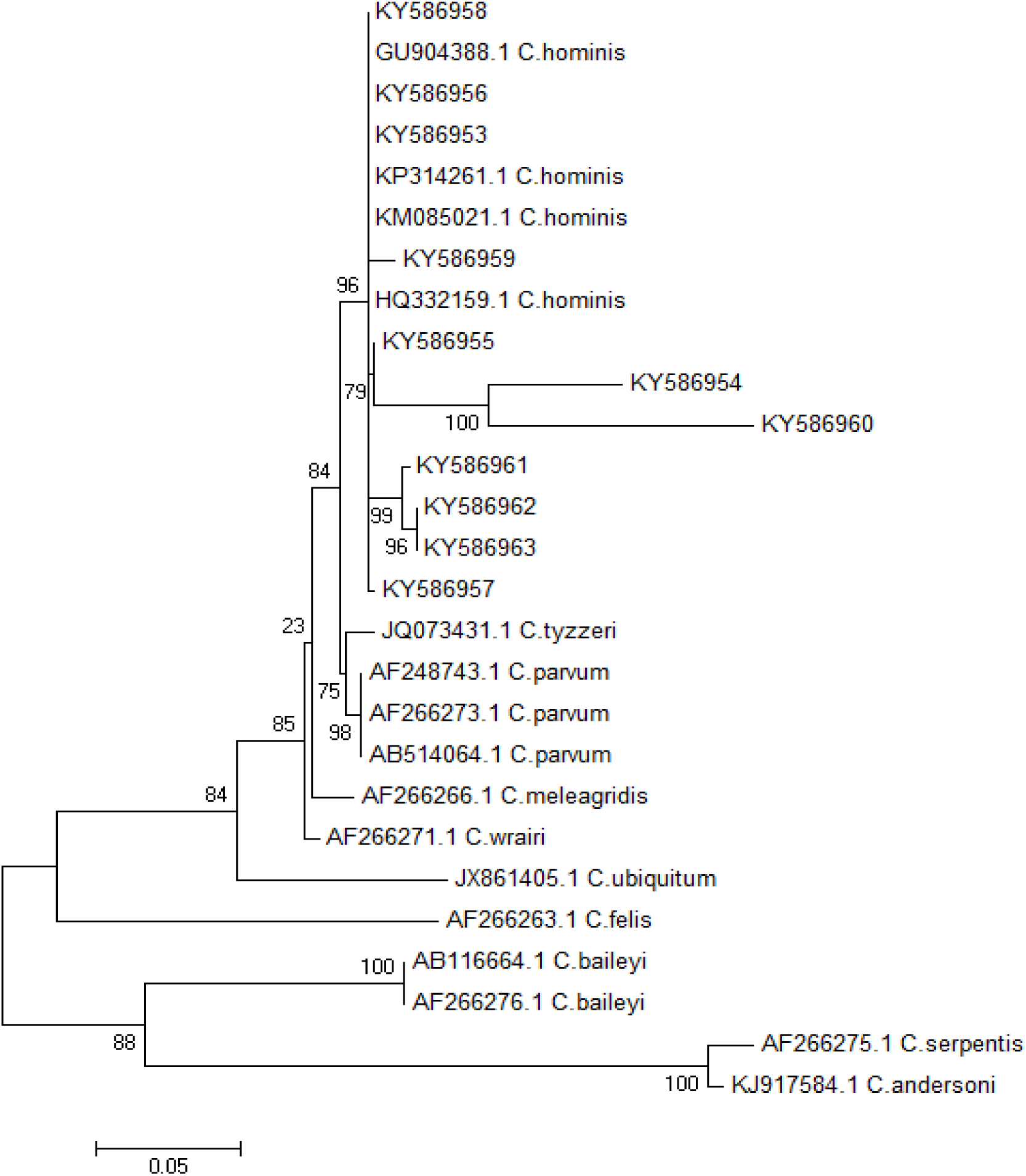
Dendogram of *Cryptosporidium* sequences isolated from cattle in Kiruhura district, south western, Uganda. The tree with the highest log likelihood (−2058.7706) is shown. The percentage of trees in which the associated taxa clustered together is shown next to the branches and is estimated from 1,000 re-samplings of the sequence data. Reference sequences are shown with GenBank accession numbers and species name. The scale bar indicates nucleotide substitutions per site.

## Discussion

The aim of this study was first to determine the prevalence of *Cryptosporidium* in cattle bordering the LMNP using microscopy. The study also aimed to genotype the isolated *Cryptosporidium* species in order to determine if the cattle posed a zoonotic threat to the local human population.

The overall *Cryptosporidium* prevalence in this study was 7.7% which is comparable to the prevalence 7.7% and 7.8% obtained in Kenya and Ethiopia by Kang’ethe *et al*., (2012) and Wegayehu, Adamu, & Petros, (2013) respectively. However the prevalence obtained in this study is higher than 2.2% previously reported in western Uganda by Salyer, Gillespie, Rwego, Chapman, & Goldberg, (2012). The possible explanation for this difference could be due to the variation in sampling techniques as well seasonality. The samples in this study were collected during the dry season when higher pressure is exerted on the scarce water sources which results poor sanitation practices that facilitate transmission (Kang’ethe *et al*., 2012). Furthermore, the prevalence reported in this study was much lower than a previous report of 38% in calves by Nizeyi *et al*., 2002. This difference in prevalence could be due to age-related susceptibility with calves at a higher risk of *Cryptosporidium* infection than adult cattle because of their naive immunological status (Brook, Hart, French, & Christley, 2008; Maddox-Hyttel, Langkjær, Enemark, & Vigre, 2006; Maikai *et al*., 2011; R. P. Smith, Cheney, & Giles, 2014).

In this study, breed associated differences in the distribution of *Cryptosporidium* infection was observed. The infection rate of *Cryptosporidium* in crosses (9.2%) was higher than that of the Friesian breed (7.1%), Ankole (5.7%) and Boran (2.8%). This variation could be due to native breeds being more resistant to diseases than the exotic breeds and crosses (Mwai, Hanotte, Kwon, & Cho, 2015). However this difference in infection was found to not be statistically significant. Age was not a risk factor in the prevalence of *Cryptosporidium* infections in this study however this differed from reports in previous studies where calves were at a significantly higher risk of infection as compared to adult cattle (Fayer *et al*., 2007).

For molecular analysis, the 553bp COWP gene product was successfully amplified in 20 of the 28 microscopy positive samples. The unsuccessful amplification of expected DNA fragment in the rest of microscopy positive samples may be explained by the low oocyst concentration in the faecal samples analysed. Furthermore, the unsuccessful amplification could have been due to fecal constituents such as bilirubin, bile salts, and complex polysaccharides that inhibit PCR (Morgan, F. U., Pallant, L., Dwyer, B. W., Forbes, D. A., Rich, G., & Thompson, 1998; Thornton & Passen, 2004). In addition the COWP gene primers in general only amplify DNA of *C. hominis, C. meleagridis C. parvum* and species or genotypes closely related to *C. parvum*. This narrow specificity may also have led to the failure to successfully amplify the 8 isolates (Xiao, 2010).

BLAST search comparison of the sequenced COWP gene fragments indicated that the all sequences generated in this study (KY586953-KY586963) are *C. hominis*. These *C. hominis* sequences however also showed great similarity with *C.parvum* and this is because there is only a 3-5% sequence divergence between *C. hominis* and *C. parvum* (Xu *et al*., 2004). Phylogenetic analysis of gene sequences from this study showed that all the sequences clustered into a single clade, with known *C. hominis* sequences from the GenBank, with a bootstrap value of 95%. This bootstrap value indicates that this clustering is highly supported and further emphasizes that the sequences generated from this study were isolated from *C. hominis*.

The findings of the present study indicate that *Cryptosporidium spp*. infections are prevalent in cattle in Kiruhura district. This is the first report documenting the isolation of *C. hominis* from cattle in Uganda. It is generally accepted that *C. hominis* primarily infects humans with no animal reservoir. However there is growing evidence indicating that *C. hominis* infects livestock (Giles *et al*., 2009; Guk, Yong, Park, Park, & Chai, 2004; Kang’ethe *et al*., 2012; Rajendran *et al*., 2011; Smith *et al*., 2005). This study further substantiates of arthroponotic transmission of *Cryptosporidium* from humans to cattle indicating that animals may play an important role in the epidemiology of human cryptosporidiosis (Giles *et al*., 2009). The role played by animals in the epidemiology of human cryptosporidiosis has been controversial, particularly as potential zoonotic reservoirs of infection. This study can hopefully contribute to this discussion.

## Conclusions

The findings of this have significant public health implication because this study represents a natural completed life cycle of *C. hominis* in the bovine host and as a result necessitates molecular epidemiological studies in order to elucidate the sources of *C. hominis* and the transmission patterns. This will provide more information in understanding the role played by cattle in human cryptosporidiosis.

## List of abbreviations

COWP: *Cryptosporidium* Oocyst Wall Protein
DVO: District Veterinary Officer
LMNP: Lake Mburo National Park
PCR: Polymerase Chain Reaction
MAAIF: Ministry of Agriculture Animal Industry and Fisheries
UBOS: Uganda Bureau of Statistics

## Declarations

### Ethics approval and consent to participate

This study was reviewed and approved by the Research Ethics Committee, College of Health Sciences, Makerere University under the reference number REC/2011/195. Written and verbal consent was obtained from the farmers before the sampling of animals.

### Availability of data and materials

The datasets used and/or analysed during the current study are available from the corresponding author on reasonable request.

### Authors’ contributions

SGW contributed to the conception of the idea, design, sample collection, data analysis and preparation of the manuscript. AK contributed to conception of the idea, study design and interpretation of results and manuscript preparation. GA contributed to sample analysis and drafting of the manuscript. CK contributed to developing the study design, interpretation of results and preparation of the manuscript. All authors read and approved the manuscript.

## Acknowledgement

Special thanks go to Phillip Kimuda Magambo, Kiseka Henry, Jonan Tubenawe, Monica Nambi and Mary Frances Nakamya for numerous helpful suggestions during the study.

